# Depleting trafficking regulator CASK promotes intercalated disc organization and ventricular function

**DOI:** 10.1101/2024.10.14.618172

**Authors:** Camille Blandin, Gilles Dilanian, Vincent Fontaine, Nathalie Mougenot, Basile Gravez, Pierre Bobin, Laetitia Duboscq-Bidot, Dounia Farhi, Solenne Chardonnet, Sophie Nadaud, Jose L. Sanchez-Alonzo, Andriy Shevchuk, Julia Gorelik, Estelle Gandjbakhch, Stéphane Hatem, Eric Villard, Elise Balse

## Abstract

**Background:** The intercalated disc (ID) electromechanically couples adjacent cardiomyocytes. Alteration of this structure plays a central role in cardiac arrhythmias, notably arrhythmogenic cardiomyopathy (ACM), an inherited genetic disorder of desmosomes. Calcium/Calmodulin-Dependent Serine Protein Kinase (CASK), a costameric component of the lateral membrane, regulates cardiomyocyte protein trafficking. Here, we investigated CASK regulation of the organization of ID components in normal and pathological contexts. Here, we investigated CASK regulation of the organization of ID components in normal and pathological contexts.

**Methods:** We studied the outcomes of CASK depletion in neonatal rat hearts using a cardiac-specific adeno-associated virus strategy. We used conventional and strain echocardiography, hemodynamics, electrocardiography, histology, and gene and protein expression studies to characterize adult rat hearts. We also studied the effects of CASK depletion in neonatal rat ventricular cardiomyocytes (NRVM), and control and ACM (*PKP2*^+/-^) IPS cell-derived CM (hIPS-CM) using proteomics, electron microscopy, high-resolution imaging, mechano-scanning ion conductance microscopy (mechano-SICM), and stress resistance tests. We studied cardiac CASK expression and localization in human ACM and non-transplantable control hearts.

**Results:** Depletion of CASK in our multiple experimental models revealed that CASK regulates cardiomyocyte IDs. In rats, CASK depletion improved contractile reserve and compliance. In cultured rat cardiomyocytes, CASK knockdown increased localization of connexin 43 (Cx43) and PKP2 at IDs, resulting in increased contact stiffness. In *PKP2^+/-^* hIPS-CMs, CASK expression was increased. CASK depletion in these cells promoted PKP2 accumulation at cell contacts, formation of desmosome-like structures, and stress resistance. In the right ventricles of ACM patients, CASK protein level was also increased and CASK abnormally localized at the ID.

**Conclusion:** CASK functions as a repressor of ID organization and tissue cohesion, suggesting novel mechanisms for regulating ID structure and function. These observations, along with CASK upregulation and mislocalization in ACM, open up new perspectives on understanding the pathophysiology of ACM and suggest innovative strategies for its treatment.

## INTRODUCTION

Cardiomyocytes are highly differentiated cells with specialized membrane domains. Costameres (i.e., focal adhesions) of the lateral membrane provide the link between the extracellular matrix and the sarcomere along the long axis of the cardiomyocyte^1^. The intercalated disc (ID), which electrically and mechanically couples adjacent cardiomyocytes termini, comprises macromolecular complexes with distinct properties: fascia adherens and desmosomes involved in cell adhesion, and gap junctions and complexes of cardiac sodium channels involved in electrical propagation.

The ID is compromised in arrhythmogenic cardiomyopathy, a condition characterized by a high propensity for life-threatening arrhythmias in young individuals and progressive adipofibrous infiltration of the ventricular walls at the expense of muscle mass^2–5^. The majority of ACM cases are caused by pathogenic variants in genes encoding desmosomal proteins, with plakophilin-2 (*PKP2)* loss-of-function variants most frequently affected^5^. The mechanisms linking desmosome gene mutations to these phenotypes remain incompletely understood.

Members of the membrane-associated guanylate kinase (MAGUK) proteins play a crucial role in the organization of cell polarity, cell-cell adhesion and the regulation of macromolecular complexes^6^. CASK (Calcium/Calmodulin-Dependent Serine Protein Kinase), a member of this superfamily, is primarily known for its key role in synapse organization in neurons^7,8^. In cardiomyocytes, we previously showed that CASK interacts with the costameric dystrophin-glycoprotein complex at the lateral membranes and regulates the trafficking and membrane targeting of the cardiac sodium channel^9,10^ at the sarcolemma. In the present study, we investigated whether CASK regulates the targeting of other ID components. We provide evidence that CASK is a crucial regulator of ID organization, regulating gap junction and PKP2 accumulation at IDs. We show that CASK is upregulated and mislocalized to IDs in ACM, suggesting a novel pathophysiological mechanism. These results provide two significant insights. First, they establish the major role of the lateral membrane in the organization of the ID. Second, they show that CASK could be targeted to mitigate the ACM phenotype.

## METHODS

Extended methods are available in the Supplemental Material.

### Animals

All studies were performed on neonatal or 2-month-old (P60) male Wistar rats (Janvier Labs, France). Experimental procedures were carried out in accordance with the French Institutional guidelines for animal experimentation (MESR, authorization #22492) and conformed to the Directive 2010/63/EU of the European Parliament.

### Human hearts

Human control (N=6, non-failing, non-transplantable hearts) and ACM (N=6, heart transplant surgery) right ventricle explants were collected. ACM samples were collected under authorizations NCT05569356, CPP 2022-A00283 40 and AC-2019-3502^11^. The control tissue samples were provided by the Biological Resources Center ICAN Biocollection (Institute of Cardiometabolism and Nutrition, Sorbonne University). For Western blots, 2 *DSG2* mutation carriers, 1 *PKP2* mutation carrier and 2 ACM samples without desmosome mutation (none) were used. For immunohistochemistry, 2 control samples and 2 ACM samples, carrying *DSG2* mutations, were used.

### Design of viral vectors

shRNA sequences for CASK 5’-tggagaatgtgaccagagttcgcctggta-3’ (shCASK) and control (shscr) 5’-gcactaccagagctaactcagatagtact-3’ were provided by Origen into a pRFP-C-RS vector. shCASK or shscr under a U6 promoter were subcloned into pShuttle vector containing an IRES-GFP reporter and produced in adenovirus (CPV, UMR1089, Nantes). ShSCR expressed a scrambled sequence of the shCASK without predicted target in the rat or human reference genomes. NRVM and hIPS-CM were treated with adenovirus at 5.10^7^ vg/ml. Adeno-associated virus (AAV) was generated using an optimized AAV2/9 backbone (custom pENN-AAV vector) containing the Tnnt2 cardiac promoter driving GFP with a mir30/mirE scaffold containing targeting sequences for CASK 5’-caggaattataatgcttatcta-3’ (AAV-shCASK) in the 3’ UTR. A scrambled sequence, 5’-cggagaatgtgaccagagttcg-3’, was used for control (AAV-shSCR). One-day-old male Wistar rats received a single jugular vein injection of AAV (8.10^10^ vg/50µl diluted in 0.9% NaCl solution).

### Echocardiographic assessment of cardiac function

Conventional and strain echocardiographic measurements were performed on lightly anesthetized rats (2-3% isoflurane and 2L/min oxygen flow rate). Body temperature was carefully monitored and maintained at 37°C (VisualSonics rat platform) during the entire procedure. Transthoracic echocardiography was carried using Vevo 3100 VisualSonics (Fujifilm), with a 15-30 MHz probe (MX250). From the parasternal long axis 2D view, LV wall thickness and cavity diameters were measured in M-mode. Isoproterenol (50 mg/kg/ml in 0.9% NaCl) was administered IP. Global longitudinal strain of the left ventricle was measured from parasternal long-axis images using speckle-tracking-based imaging. Recordings were quantified and averaged for three cardiac cycles. Acquistion and analysis were performed by an operator blinded to the rat groups.

### Invasive hemodynamic measurements

8-week-old Wistar male rats were anesthetized with a mixture of ketamine/xylazine (100 mg/kg/10 mg/kg, i.p.). Animals were placed on controlled heating pads, and the temperature was maintained at 37°C. Invasive hemodynamic measurements were then performed on spontaneously breathing rats using a pressure transducer catheter (SPR-838, 2F Millar Mikro-tip P-V catheter, Millar Instruments) introduced into the right carotid artery and advanced into the ascending aorta. After stabilization for 5 min, arterial blood pressure was recorded. The catheter was advanced into the LV under pressure control. After stabilization for 5 min, signals were continuously recorded at a sampling rate of 1000 samples/s using a P-V conductance system (MVPS-Ultra, Millar Instruments) connected to a PowerLab 16/30 data acquisition system (AD Instruments), stored, and displayed by the Lab-Chart 5 Software System (AD Instruments). LV P-V relations were also registered during transient compression of the inferior vena cava (reducing preload) using a syringe pusher. Indices of LV contractility (ESPVR and PRSW) and compliance (EDPVR) were analyzed using P-V analysis program (PVAN; Millar Instruments).

### Isolation and culture of rat cardiomyocytes

Adult ventricular cardiomyocytes were obtained by enzymatic dissociation on a Langendorff column as previously described^12^ and used for immunofluorescence immediately after isolation. Neonatal ventricular cardiomyocytes (NRVM) were obtained by enzymatic dissociation of 1-day-old Wistar rats heart using Neonatal Heart Dissociation Kit (Miltenyi Biotec). Ventricles were cut into 1-2 mm pieces and incubated with digestive enzymes (37°C, 5% CO_2_) and myocytes were mechanically extracted using the gentleMACS^TM^ dissociator. Cells were cultured under standards conditions (37°C, 5% CO2) in DMEM medium (Gibco) supplemented with 1% penicillin/streptomycin (Gibco) and 10% fetal bovine serum (Eurobio) and used for experiments 48h after transduction.

### Cardiomyocytes derived from induced pluripotent stem cells

The *PKP2*^+/-^ cell line was generated as published^13^ using a CRISPR-Cas9 based approach from the control ICAN403.3 hiPSC line previously described^14^. hIPS-CM were plated on matrigel-coated dishes and cultured in RPMI1640+B27 medium (Life Technologies). Cardiomyocyte differentiation was assessed by flow cytometry and immunostaining as previously described^13,14^. Cells were used 48h after transduction with adenovirus.

### Mechano scanning ion conductance microscopy

The hopping probe variant of scanning ion conductance microscopy (SICM) was used to scan live cells as previously described^15^. Briefly, SICM is a scanning probe microscopy technique that uses glass nanopipettes, similar to micropipettes used for electrophysiological recordings but smaller diameter, as cell surface proximity sensor and enables non-contact 3D topographical imaging of living cells in physiological buffer mediums. Topography and stiffness mapping experiments were performed on NRVM as previously described^16,17^. The principle of stiffness mapping is based on the phenomenon of non-contact low-stress exertion when SICM pipettes with diameters less than 100 nm are used at set points higher than 1% of ion current drop from bulk.

### LC-MS/MS proteomic analysis

Protein lysates from 5 independent NRVM cultures were digested with trypsin on S-trap columns (Protifi) and analyzed by LC-MS/MS on a timsTOF Pro mass spectrometer (Bruker) coupled to a nanoElute UPLC (Bruker). The peptides were separated using a 70 min gradient (from 2 to 30 % ACN in 0.1% FA) and analysed in DDA-PASEF mode. Data analysis was performed with MaxQuant with the UniProt *Rattus norvegicus* protein database and ProStar zero was used for data filtering and statistical analysis (Limma test corrected with the Adaptive Benjamini and Hochberg procedure, FDR < 1%).

### Statistics

Gaussian distribution was tested by the Shapiro-Wilk test. Normally distributed data were compared using an unpaired t-test or one-way ANOVA, followed by a Tukey test for multiple comparisons. Data with non-Gaussian distribution were compared using the Mann-Whitney test. *P* values were considered significant at *P*<0.05. Data were prepared using Prism 8.0 software (GraphPad) and presented as means±SD.

## RESULTS

### Cardiac-specific AAV-mediated CASK silencing in rat induces physiological hypertrophy

We generated AAV9-Tnnt2-GFP-shCASK (AAV-shCASK) to express GFP and shRNA targeting CASK specifically in rat cardiomyocytes. AAV9-Tnnt-GFP-shScr (AAV-shScr), expressing a scrambled shRNA, was used as control. AAV was delivered at postnatal day 1 (P1) (Figure 1A). Although AAV expressed the transgene in a mosaic distribution (Supplemental Figure 1A), western blot on total heart lysate showed a significant, 67% reduction in CASK expression two months after injection (Supplemental Figure 1B). Decrease in CASK expression was also observed by fluorescence quantification on freshly isolated GFP^+^ cardiomyocytes (Supplemental Figure 1C). AAV-shCASK hearts had increased width but no difference in the heart weight to tibia length ratio compared to controls (Figure 1C). Echocardiography revealed thinning of the heart walls associated with an increase in left ventricular end-diastolic diameter (Figure 1D). An increase in cardiac output, dependent of increased stroke volume, was also observed (Figure 1E). CASK depletion reduced ejection fraction in the basal state (Figure 1F). However, adrenergic stimulation (isoproterenol, ISO) showed that contractile reserve was similar to that of control hearts (Figure 1F). In addition, pressure-volume recordings revealed that CASK-depleted hearts had elevated enhanced contractility (ESPVR and PRSW) and compliance (EDPVR) (Figure 1F). This result was consistent with assessment of contractility by echocardiographic speckle-tracking (left ventricle global longitudinal strain, Figure 1G). No electrophysiological alterations were observed on ECG (Supplemental Table 1B), even when rats were challenged with flecainide (Supplemental Table 1C). Tissue analysis by TUNEL staining or Bax expression by RT-qPCR revealed no change in apoptosis (Supplemental Figure 1D-E). Masson’s trichrome staining showed no evidence of fibrosis (Supplemental Figure 1F). This result was consistent with assessment of contractility by echocardiographic speckle-tracking (left ventricle global longitudinal strain, Figure 1G). No electrophysiological alterations were observed on ECG (Supplemental Table 1B), even when rats were challenged with flecainide (Supplemental Table 1C). Tissue analysis by TUNEL staining or Bax expression by RT-qPCR revealed no change in apoptosis (Supplemental Figure 1D-E). Masson’s trichrome staining showed no evidence of fibrosis (Supplemental Figure 1F). In addition, RT-qPCR of mRNA expression of basement membrane components, metalloproteinases, or metallopeptidase inhibitors showed no differences between AAV-shCASK and AAV-shSCR hearts (Supplemental Figure 1G), supporting no remodeling of extracellular matrix. Finally, no difference was observed in the MYH7/MYH6 ratio or the BNP expression level (Supplemental Figure 1H-I). These data show that AAV-shCASK successfully depleted CASK, and that CASK depletion did not impair and even enhanced cardiac contractility and relaxation and did not induce markers of cardiomyocyte pathological stress.

**Figure 1.**
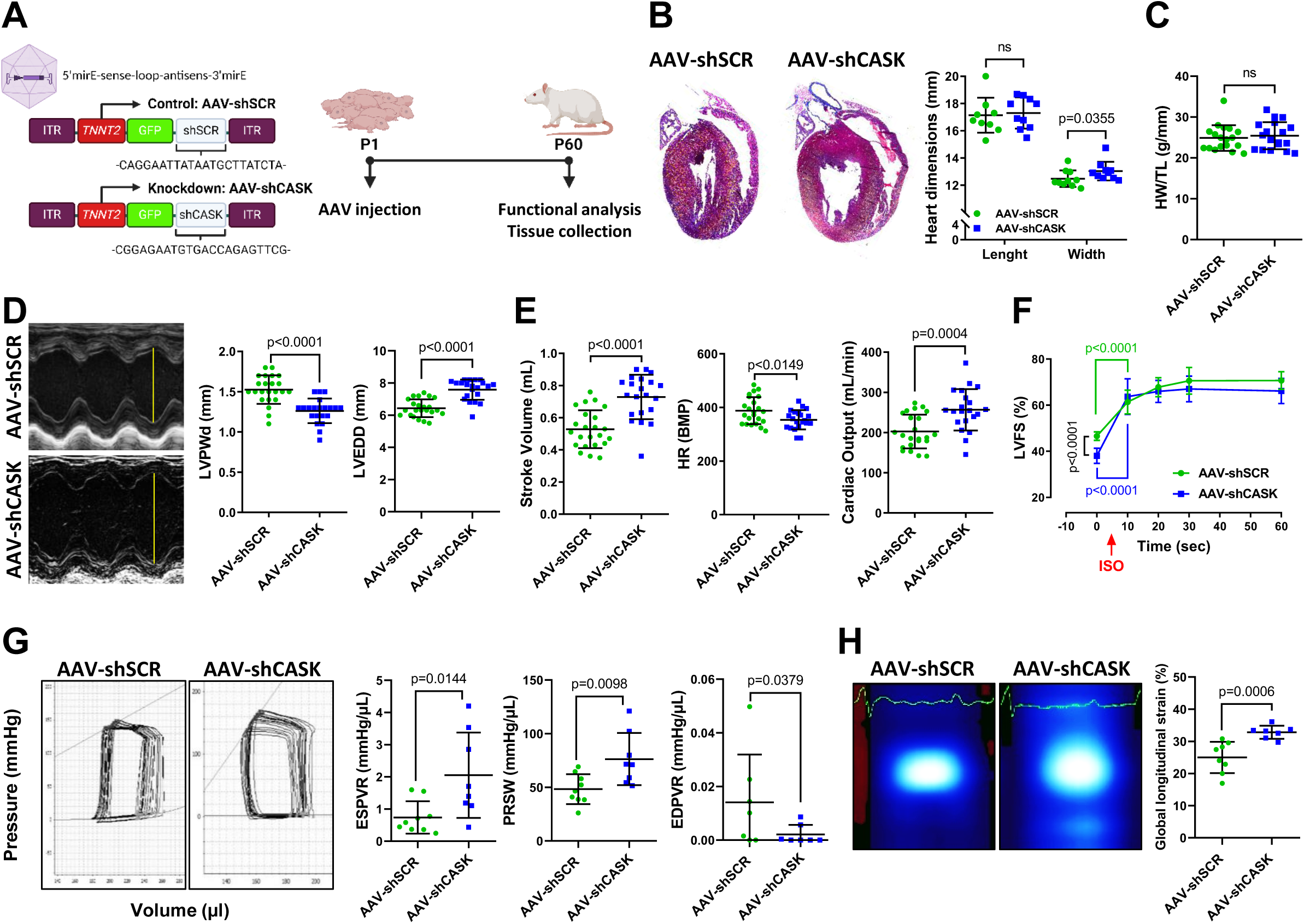
Cardiac-specific CASK silencing in rat induces hypertrophic remodeling and increased cardiac function. **(A)** Design of adeno-associated virus constructs and experimental protocol. **(B)** Representative images of Masson’s trichrome staining of AAV-shSCR or CASK-invalidated (AAV-shCASK) hearts and quantification of macroscopic parameters (N=10 rats/group, mean±SD, Mann-Whitney test). **(C)** Heart weight/tibia length ratio (N=18 rats/group, mean±SD, t-test). **(D)** Representative images of M-mode echocardiography of left ventricle, and quantification of posterior wall thickness in diastole and left ventricular end-diastolic diameter. **(E)** Cardiac output (N=21 to 23 rats/group, mean±SD, t-test). **(F)** Left ventricular ejection fraction and left ventricular fractional shortening kinetics before (t_0_) and following β-adrenergic (ISO) stimulation (N=11 to 13 rats/group, mean±SD, 2-ways ANOVA). **(G)** Representative pressure-volume loops of control and CASK-invalidated rats and quantification of systolic function parameters (N=7 to 9 rats/group, mean±SD, t-test; end systolic pressure volume relationship (ESPVR), preload recruitable stroke work (PRSW), and end diastolic pressure volume relationship (EDPVR). **(H)** Representative 2D images of ventricular deformation and quantification of global longitudinal strain (N=7 to 8 rats/group, mean±SD, t-test).

### CASK regulates the cytoskeleton and the organization of cell-cell contacts in cardiomyocytes

To further investigate the molecular consequences of CASK invalidation in cardiomyocytes, we utilized LC-MS/MS proteomics in a cell culture model of confluent neonatal rat ventricular cardiomyocytes (NRVM). CASK was depleted by 76% using adenoviral vector (Ad-shCASK). Scrambled vector (Ad-shSCR) was used as control (Figure 2A). Proteomic analysis revealed high within group correlation and effective between-group separation (Supplemental Figure 2). A total of 3565 known proteins from the UniProt reference database were measured and tested for differential expression. Stringent threshold criteria (FDR<1%, |Fold-change| > 1.3) yielded 654 differentially expressed proteins, with 322 proteins down-regulated and 332 up-regulated in Ad-shCASK compared to control (Figure 2B). Ingenuity Pathway functional network Analysis (IPA) of the 654 differential proteins identified Rho-dependent cytoskeletal organization as the main upregulated pathway. Additional upregulated pathways included cell-matrix-related signaling (“integrin signaling”) and cell-cell contact signaling (“epithelial adhesion junction signaling”) (Figure 2C). Metascape analysis identified “actin filament-based process” and “signaling by Rho GTPases” as highly regulated pathways (Figure 2D). Both Metascape and ClueGo (Cytoscape) also highlighted “protein localization to membrane” and “membrane trafficking” as CASK-regulated pathways (Figure 2D, Supplemental Figure 3). In summary, CASK knockdown in NRVM affected the regulation of the cytoskeleton, intracellular transport, and cell membrane organization. These findings suggest that CASK may play a role in regulating cardiomyocyte polarity.

**Figure 2.**
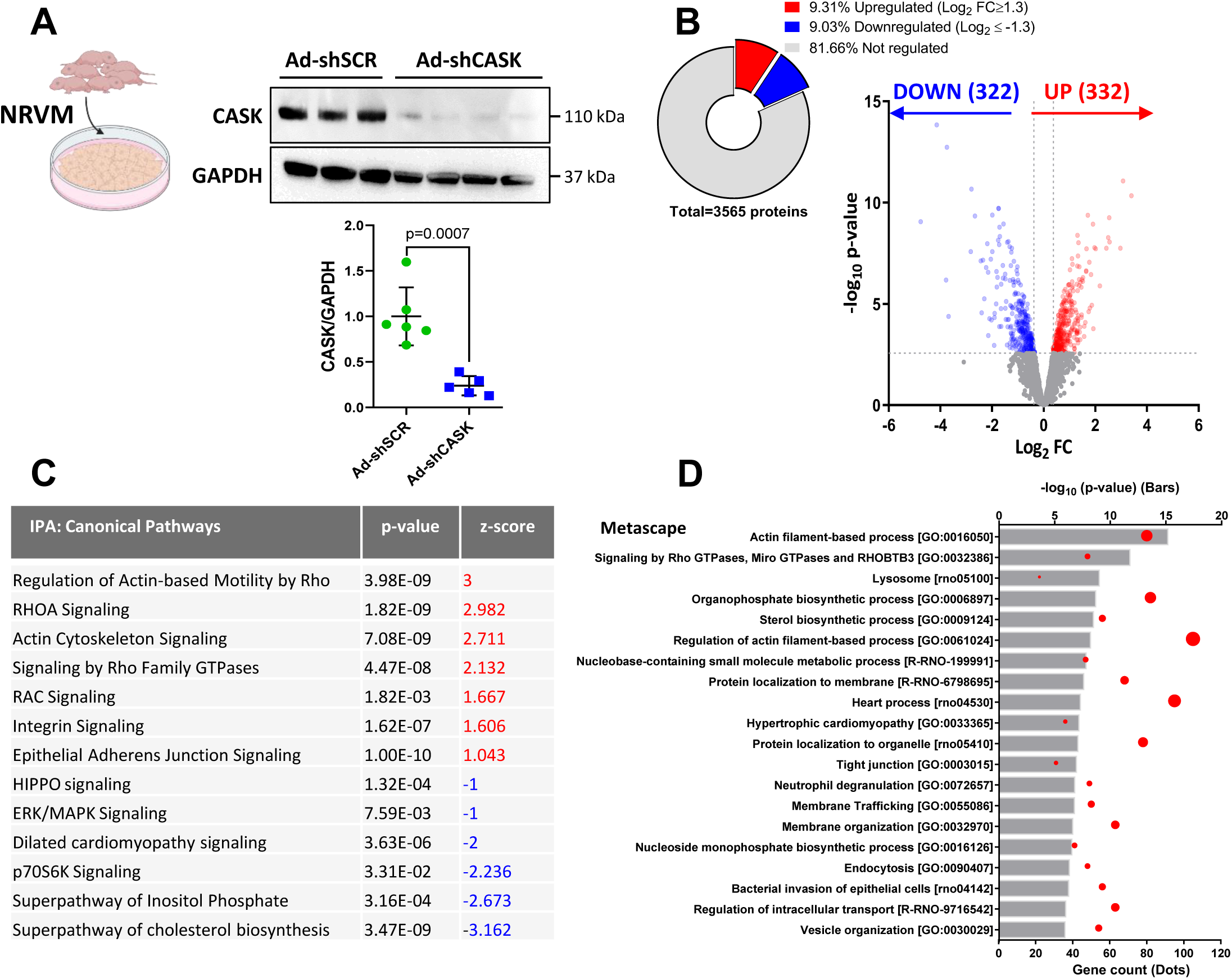
Proteomics analysis of CASK invalidation in neonatal rat ventricular myocytes (NRVM). **(A)** Representative immunoblots of CASK expression following transduction with either control (Ad-shSCR) or short hairpin RNA targeting CASK (Ad-shCASK) adenoviral constructs and quantification (N=5 to 6 hearts/group, t-test). **(B)** Volcano plot showing differentially expressed proteins between Ad-shSCR- and Ad-shCASK-NRVM (N=5 independent culture/group, FC ≥1.3 (up-regulated, red) or FC ≤1.3 (down-regulated, blue), p-value cut off 0.0027, corresponding to FDR q-value<0.01). **(C)** Differentially upregulated (red) and downregulated (blue) canonical pathways identified by Ingenuity Pathway Analysis (IPA) in CASK-silenced NRVM. **(D)** Metascape analysis showing the pathways significantly regulated following CASK invalidation. Bars indicate significance, and dots the number of genes per pathway.

### CASK-knockdown increases connexin 43 and plakophilin 2 localization at cell junctions in both NRVM and hIPS-CM

As we previously showed that CASK regulates the membrane localization of the sodium channel Na_V_1.5^9,10^, a protein highly concentrated at the ID in the cardiomyocyte^18,19^, we investigated CASK regulation of the localization of other ID proteins. We used machine-learning based image analysis to quantify contact zones from immunofluorescence images of gap junction (connexin 43, Cx43) and desmosome (plakophilin 2, PKP2) components of IDs. CASK depletion both increased Cx43 protein level (Figure 3A) and localization at contacts in NRVMs (Figure 3B). Unlike Cx43, CASK depletion reduced PKP2 protein level yet increased its localization at contacts (Figures 3A-B). We next used non-contact nanoscale mechanosensing ion conductance microscopy (mechanoSICM)^20^ to determine whether the increase in ID proteins at contacts was accompanied by changes in their mechanical properties (Figure 3C). Indeed, mechanoSICM revealed that junction height (Figure 3D, Supplemental Figure 5A) and Young’s Modulus (Figure 3E, Supplemental Figure 5B) were higher in CASK-depleted cells, confirming that CASK modified contact thickness and rigidity in cultured cells. Taken together, these data suggest that CASK is a key regulator of ID organization and governs the accumulation of ID components at contacts. To verify that CASK regulation affects protein ID localization in a human context, we repeated these experiments on cardiomyocytes derived from human induced pluripotent stem cells (hIPS-CM)^14^. Adenoviral transduction efficiency in hIPS-CM was comparable to that of NRVM (79% CASK depletion, Figure 3F vs Figure 2A). In the hIPS-CM model, Cx43 and PKP2 expression levels were not altered by CASK depletion (Figure 3G). However their localization at contacts was strongly increased (Figure 3H). We hypothesized that CASK inhibition promotes anterograde protein trafficking to the plasma membrane. To test this, we used Brefeldin-A (BFA) to prevent ER to Golgi trafficking following adenoviral transduction. In CASK-depleted hIPS-CM, BFA treatment prevented Cx43 and PKP2 accumulation at cell junctions (Figure 3I). These observations suggest that the effect of CASK depletion on ID protein localization is mediated by early anterograde trafficking of gap (Cx43) and desmosome (PKP2) proteins, rather than by altered protein level. These studies demonstrated that CASK regulates the organization of ID components, and its depletion enhanced the organization of ID components *in vitro* in both human and rat cardiomyocytes.

**Figure 3.**
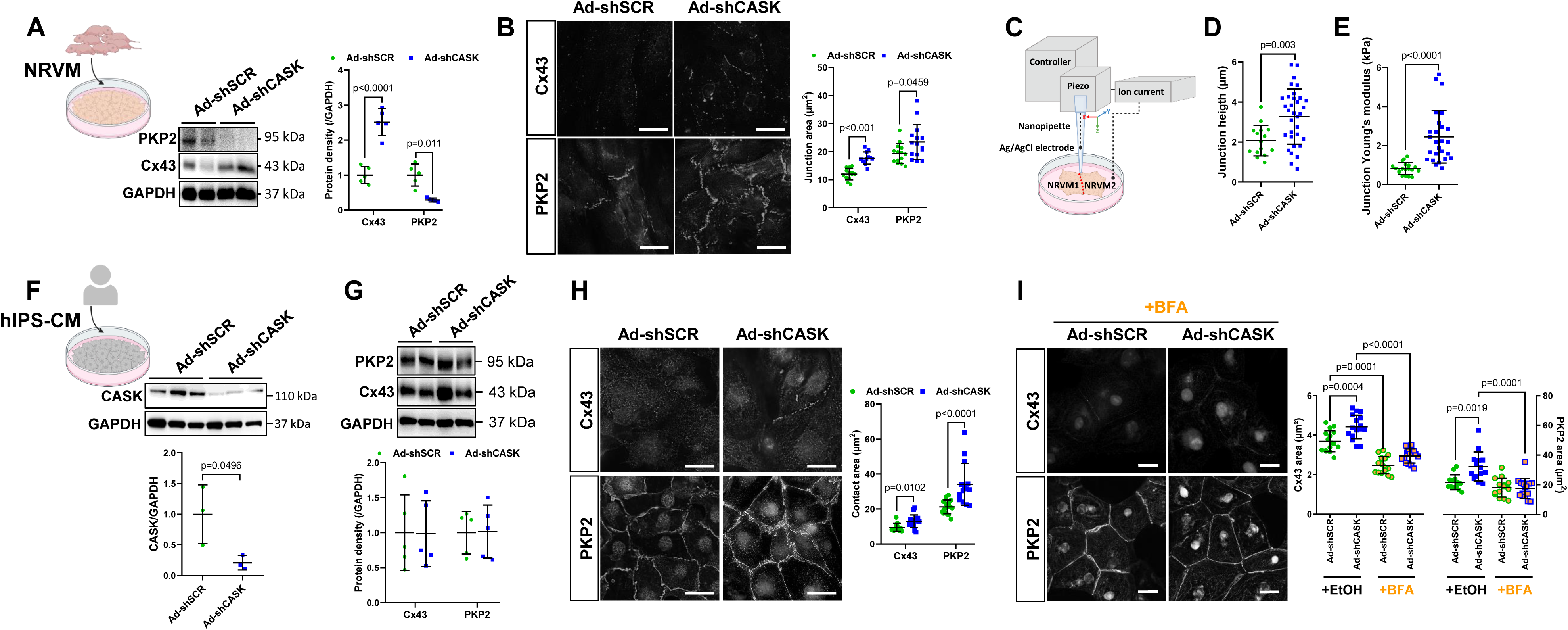
NRVM silenced for CASK exhibit cell contacts remodeling. **(A)** Representative immunoblots of Cx43 and PKP2 obtained from lysates used for LC-MS/MS experiments and quantification (N=5/group, mean±SD, t-test). **(B)** Representative immunostainings of Cx43 (top) and PKP2 (bottom) and quantification of the contact area in control and CASK-invalidated confluent NRVM (N=3 independent cell cultures, n=13 to 16 fields/culture, mean±SD, t-test, scale bar 20 µm). **(C)** Schematic diagram of mechano-SCIM directed at contacts in NRVM (for the sake of clarity, only two myocytes are represented with the junction site highlighted in red). **(D)** Quantification of junction height (N=2 independent cultures, n=15 to 33 recordings, mean±SD, t-test). **(E)** Quantification of Young’s modulus at junctions in control and CASK-invalidated NRVM (N=2 independent cultures, n=19 to 27 recordings, mean±SD, t-test). **(F)** Representative immunoblots and corresponding quantification of CASK expression in hIPS-CM showing the transduction efficiency of the shCASK adenovirus (N=3 independent differentiations/condition, mean±SD, t-test). **(G)** Representative immunoblots of PKP2 and Cx43 in whole lysates from hIPS-CM and corresponding quantification (N=5 independent differentiations/condition, mean±SD, t-test). **(H)** Representative immunofluorescence images of Cx43 (top) and PKP2 (bottom) stainings of hIPS-CM transduced with either Ad-shSCR or Ad-shCASK and quantification of contact areas. **(I)** Representative immunofluorescence images of Cx43 (top) and PKP2 (bottom) stainings of BFA-treated hIPS-CM transduced with either Ad-shSCR or Ad-shCASK, and quantification of contact areas (EtOH served as vehicle for BFA, N=2 independent differentiations/condition, n=7 to 8 field/differentiation, mean±SD, t-test, scale bar 20 µm).

### CASK invalidation in a human *in vitro* ACM model restores desmosomal integrity

Since CASK depletion upregulates PKP2, a key desmosome component whose haploinsufficiency often causes ACM^2–5^, we explored whether CASK knockdown could promote desmosome organization in this disorder. To test this hypothesis, we used an hIPS-CM cell line carrying an ACM-causing heterozygous early stop mutation of *PKP2* (*PKP2*^+/-^ hIPS-CM) that we previously generated^13^ (Supplemental Figure 6A). These cells expressed half the normal level of PKP2, mimicking the haploinsufficiency state of the disease (Figure 4A, Supplementary Figure 6B). CASK expression levels were 6-fold compared to the parental, isogenic control hIPS-CM line, indicating a potential functional link between the two proteins (Figure 4A). *PKP2*^+/-^ hIPS-CMs also were much larger than control hIPS-CMs (Supplemental Figure 6C). CASK depletion reduced the size of *PKP2*^+/-^ cells (Supplemental Figure 6E). Remarkably, CASK depletion significantly increased PKP2 localization at contacts in *PKP2^+/-^* hIPS-CM (Figure 4B) without significantly affecting total PKP2 protein levels (Figure 4C). Ultrastructural analysis of cell contacts by TEM revealed the presence of desmosome-like structures only in CASK-depleted *PKP2*^+/*-*^ hIPS-CMs, as well as smaller gaps between adjacent cardiomyocytes (Figure 4D, Supplemental Figure 6D). We therefore tested whether depleting CASK could functionally improve contact stability in *PKP2*^+/-^ hIPS-CMs using a battery of tests that measure cell-cell adhesion. CASK depletion reduced monolayer fragmentation induced by dispase treatment and shaking (Figure 4E). CASK depletion also reduced *PKP2*^+/-^ hIPS-CM cell detect caudes by detachment and an increase in live cells (Figure 4F). Finally, CASK invalidation promoted resistance to uniaxial stretching in the *PKP2*^+/-^ hIPS-CM, quantified by the number of holes formed under stretching at different time points (Figure 4G). Collectively, these results demonstrate that CASK depletion significantly attenuates the ACM phenotype of *PKP2*^+/-^ hiPSC-CMs by reducing cell hypertrophy, promoting PKP2 localization at contacts and desmosome organization, and improving the resistance of desmosomal contacts to stress.

**Figure 4.**
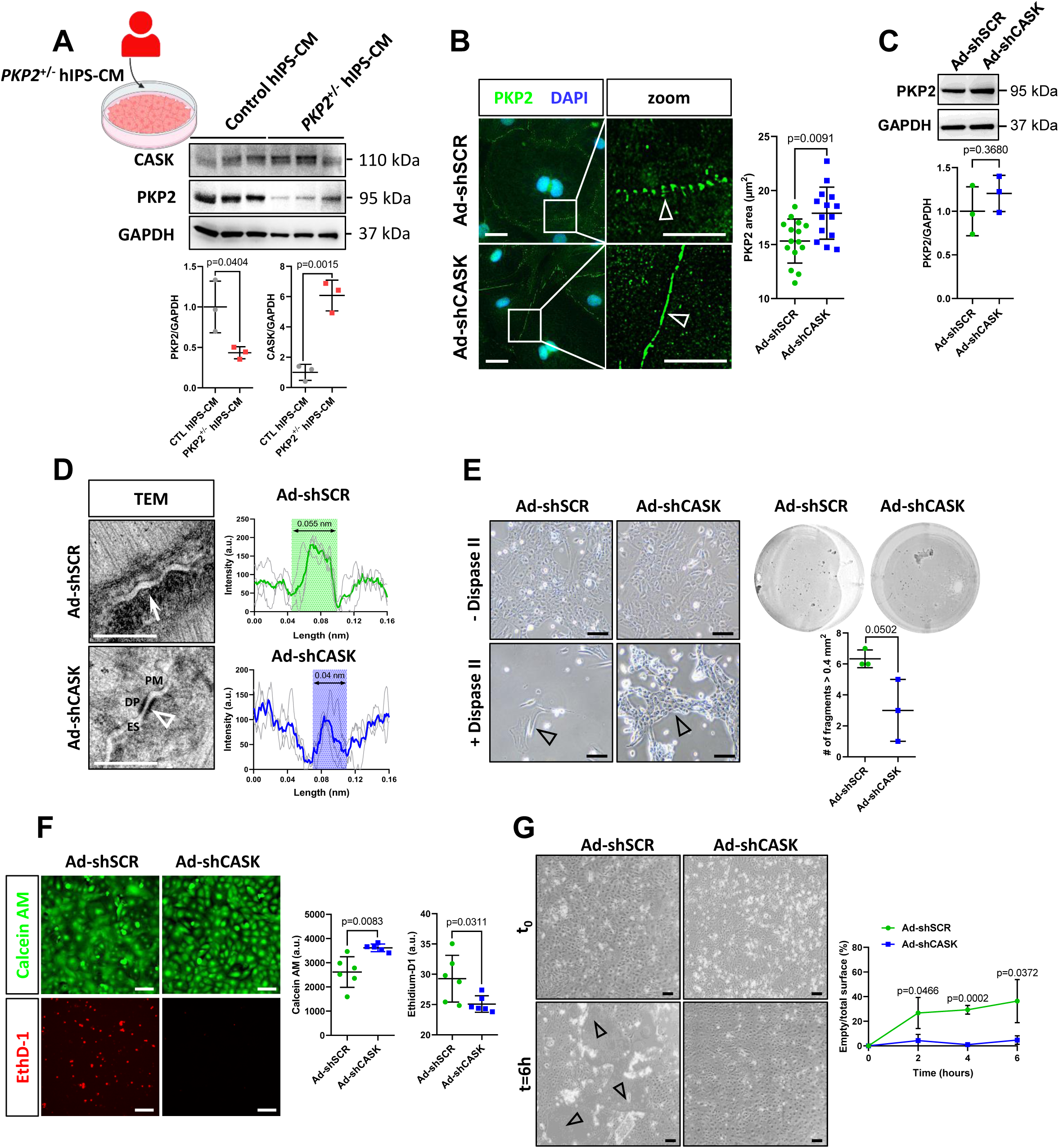
CASK silencing in PKP2^+/-^ hIPS-CM restores desmosomal integrity. **(A)** Representative immunoblots of CASK and PKP2 protein levels in control hIPS-CM and *PKP2*^+/-^ hIPS-CM (N=3 differentiations/condition, mean±SD, t-test). **(B)** Representative immunostainings of PKP2 in *PKP2*^+/-^ hIPS-CM transduced with either Ad-shSCR or Ad-shCASK and quantification of the PKP2 contact area (N=3 independent differentiations/condition, n=15 fields/differentiation, mean±SD, t-test, scale bar 20 µm, 5 µm on the zoom). **(C)** Representative Western blots of PKP2 protein in *PKP2*^+/-^ hIPS-CM showing that CASK did not increase PKP2 protein levels (N=3 differentiations, mean±SD, t-test). **(D)** Representative transmission electron microscopy (TEM) images of contact ultrastructure in control and CASK-silenced *PKP2*^+/-^ hIPS-CM, and representation of the average distance between two adjacent cells (N=1 differentiation, n=3 images/condition, scale bar 500 nm). White arrows highlight fascia adherens, white arrowheads highlight desmosome-type contacts. DP: dense plaque, ES: extracellular space, PM: plasma membrane. **(E)** (left) Representative bright field images of the *PKP2*^+/-^ hIPS-CM monolayer transduced with either Ad-shSCR or Ad-shCASK and submitted to enzymatic (+ Dispase II) or not (-Dispase II) and mechanical dissociation. Arrow heads highlight remaining contacts in PKP2^+/-^ hIPS-CM invalidated for CASK (scale bar 100 µm). (right) Representative bright filed images of full 35 mm^2^ dishes of *PKP2*^+/-^ hIPS-CM after the dispase assay and quantification of the number of fragments (>4mm^2^) after shaking-induced monolayer dissociation (N=1 differentiation, n=3 dishes/condition, mean±SD, t-test). **(F)** Anoïkis assay and quantification of live cells (Calcein AM) and dead cells (Ethidium-D1) (N=2 independent differentiations, n=3 dishes/differentiation, mean±SD, t-test, scale bar 100 µm). **(G)** Representative bright filed images of PKP2^+/-^ hIPS-CM transduced with either Ad-shSCR or Ad-shCASK before (t_0_) and after being subjected to mechanical longitudinal stretching for 6 hours, and quantification of stress resistance expressed as a function of detached surface area over total well surface area at different time points (N=3 independent differentiations, mean±SD, t-test, scale bar 1 cm).

### CASK expression os increased in ACM patients

Since native levels of CASK seem to negatively regulate desmosome organization, we aimed at investigating whether its expression was altered in RV myocardium of ACM patients carrying different pathogenic variants^11^, compared to unused transplant donor controls (Supplemental Table 2). Compared to control, ACM tissues exhibited cardiomyocyte dystrophy as shown by quantification of cell area in cross sections (Figure 5A) and reduced cardiac troponin T (cTnT) levels (Figure 5B) probably due to the loss of cardiomyocytes in these advanced-stage ACM samples. Therefore, to quantify CASK specifically in the cardiomyocyte fraction, we normalized CASK and cTnT levels to total protein levels and then normalized CASK to cTnT. The ratio of CASK to TNT was greater in ACM compared to control (Figure 5B), consistent with our observations in *PKP2*^+/-^ hIPS-CMs (Figure 4A). To confirm this finding, we quantified immunofluorescently stained right ventricle sections of control and ACM patients with *DSG2* pathogenic variants. Dystrophin was employed to specifically outline the lateral membrane in tissue sections, creating a mask. Quantification of CASK signal within the dystrophin mask revealed an increase in CASK at the lateral membrane of cardiomyocytes in ACM samples (Figure 5C, Supplemental Figure 7). In addition, a marked localization of CASK at the ID was observed in ACM, a feature rarely observed in control patients (Figure 5D) and never observed in murine control tissue^9^. In conclusion, these data show that CASK was upregulated and abnormally located in patients with ACM.

**Figure 5.**
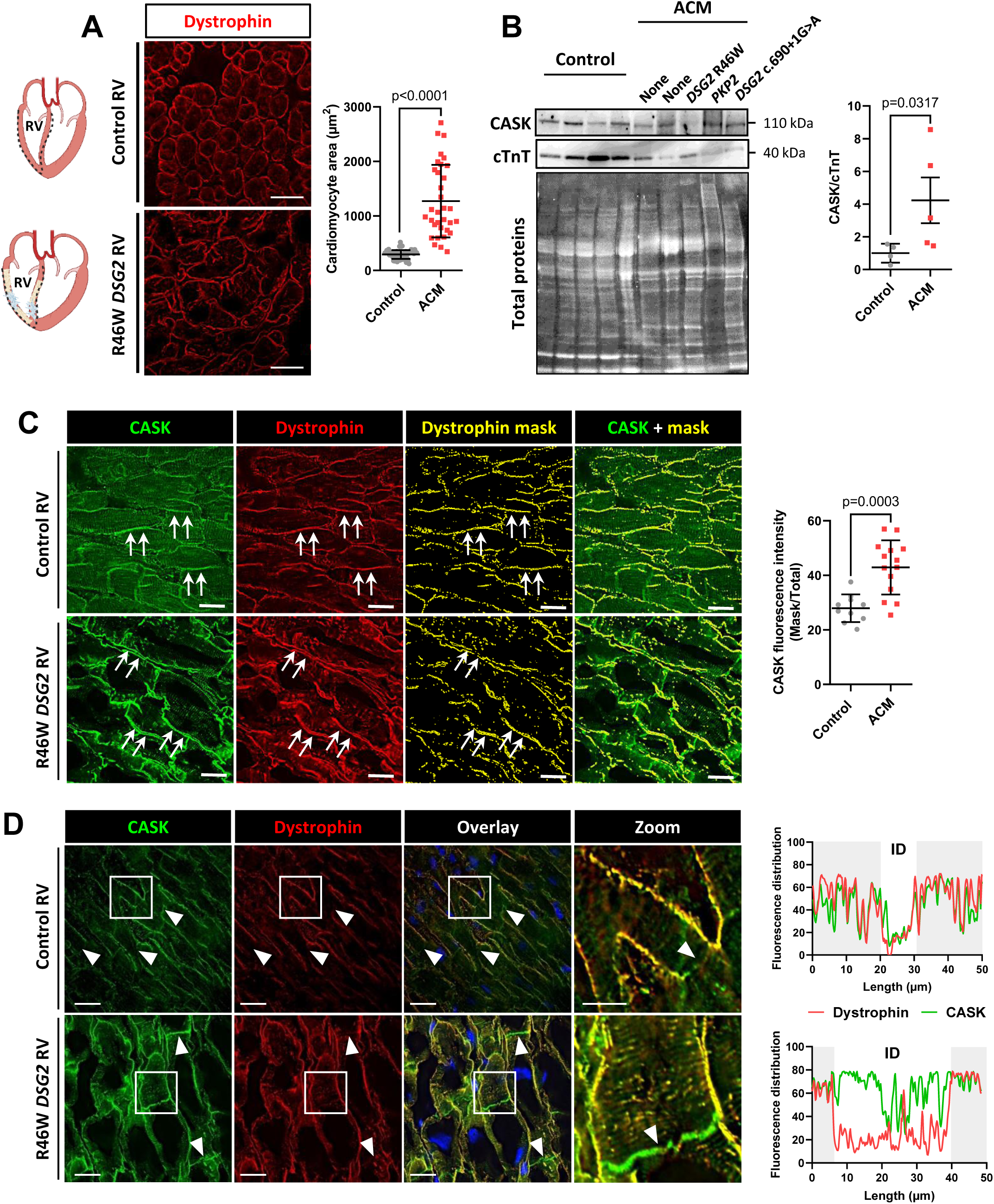
CASK expression increases in right ventricular human ACM explants. **(A)** Representative immunofluorescence images of the right ventricle of a control patient and an ACM patient carrying a mutation in desmoglein 2 (*DSG2*) and quantification of cardiomyocyte surface area measured on cross-sections with the dystrophin marker (N=2 control explants, N=3 ACM explants, n=33 to 59 fields, mean±SD, t-test, scale bar 20 µm). **(B)** Representative immunoblots of CASK and cardiac troponin T (cTNT) and corresponding total proteins obtained from right ventricle samples from control and ACM patients, and quantification of CASK expression normalized to cTNT (N=4 control explants, N=5 ACM explants, mean±SD, Mann-Whitney test). **(C)** Quantification of CASK fluorescence intensity (green) at the lateral membrane in longitudinal sections of control and ACM right ventricles using the dystrophin staining (red) to create a mask (yellow) (N=2 control explants, N=3 ACM explants, n=10 to 14 fields, mean±SD, t-test, scale bar 20 µm). Arrows point the lateral membrane. **(D)** Representative immunostainings of CASK (green) and dystrophin (red) longitudinal sections of control and ACM right ventricles (scale bar 50 µm, arrowheads highlight ID), and enlargement of white-framed areas and representation of the fluorescence distribution of the CASK (green) and dystrophin (red) along a 50 µm segment drawn on the cardiomyocyte membrane, with the ID region highlighted in white and the lateral membrane in highlighted in gray (scale bar 10 µm).

## DISCUSSION

In this study, we report for the first time that the CASK protein is involved in the organization of the ID. Our results showed that depletion of CASK in vitro increased the targeting of components of two ID structures, gap junctions and desmosomes, to cell-cell contacts. Moreover, we observed that CASK inhibition enhanced desmosome organization in a human model of ACM. In vivo, cardiac-specific CASK depletion resulted in physiological cardiac remodeling, i.e. increased systolic function and compliance without upregulation of pathological cardiomyocyte stress markers. In biopsies from ACM patients and in the human ACM model, we also observed increased CASK expression, suggesting a potential causal role of CASK is the disease.

The lateral membrane and the intercalated disc are considered to be separate entities, with no functional interplay. In the adult myocardium, CASK expression is restricted to the lateral membrane, where it associates with the dystrophin-glycoprotein complex at the costamere^9,10^, the transverse structures in the sarcolemma that align with the Z-lines of the sarcomere. Linking the extracellular matrix (ECM) to the actin cytoskeleton via transmembrane and submembrane proteins, costameres function as focal adhesions and are a crucial hub of cellular mechanosensing^1^. Mechanosensing involves feedback loops where mechanical cues activate Rho GTPases, which in turn regulate actin dynamics to modulate the cellular response^21,22^. A key finding of the present study is that the increased targeting of Cx43 and PKP2 to IDs upon CASK depletion correlated with changes in junctional stiffness, but did not correlate with total Cx43 and PKP2 protein levels, ruling out direct gene regulation of ID components. Two main sets of experiments support the idea that the effect of CASK was mediated by protein addressing to the ID. On the one hand, the proteomic study identified the actin cytoskeleton, Rho GTPase signaling and intracellular transport as being particularly affected by CASK depletion. On the other hand, PKP2 and Cx43 accumulation at the ID upon CASK depletion was prevented by blocking transport out of the Golgi with Brefeldin-A. While the precise mechanism by which CASK controls the addressing of ID proteins has not yet been fully identified, our mechanistic hypothesis, based on the mechanobiome concept^23^, is that lateral membrane remodeling impacts the actomyosin cytoskeleton. This, in turn, redistributes tensions within the cardiomyocyte, and functionally strengthen cell-cell junctions independent of gene regulation.

In ACM, several studies have reported an overall decline in desmosomal proteins, independent of mutations associated with changes in mRNA levels^24–30^. A potential mechanistic explanation for the pathophysiology of ACM was provided by a recent publication showing aberrant ubiquitination of lysine residues in PKP2, JUP and DSP proteins and that prevention of protein degradation by blocking the ubiquitin-proteasome system increased desmosomal protein levels in a mouse harboring a *Pkp2* nonsense mutation^30^. In the present study, PKP2 expression was halved in the human ACM model (*PKP2^+/-^* hIPS-CM), and CASK depletion failed to increase PKP2 expression. Nevertheless, CASK depletion strongly increased PKP2 localization at contacts, reduced intercellular space at the ID, and functionally promoted desmosome adhesion. Based on our results showing that CASK effects involved regulation of anterograde trafficking, increased CASK levels would have deleterious consequences on ID organization and stabilization. Interestingly, more abundant levels of CASK were observed in PKP2^+/-^ hIPS-CM and in human myocardium from ACM patients carrying pathogenic variants in different genes. These observations further support a potential functional link between desmosomal proteins and the lateral membrane protein CASK.

A major limitation of our study is that it relies mainly on in vitro models that do not adequately reproduce the complex three-dimensional organization of the myocardium. Although hIPS-CMs are often considered immature, notably due to the absence of a true desmosome, they offer the advantage of being able to study the effect of desmosomal mutations identified in patients (e.g.^30,31^). In the *PKP2^+/-^* hIPS-CM model used in this study, ultrastructural microscopy revealed the presence of desmosome-like structures after CASK depletion, and cell adhesion assays confirmed a beneficial functional effect. It is therefore tempting to speculate that cardiac-specific AAV-mediated partial invalidation of CASK could prove a potential therapeutic target for desmosome stabilization in ACM. Given the in vivo results showing that CASK depletion was not deleterious in rats, but on the contrary improved cardiac function, the next step will be to establish the actual functional benefits of this strategy in well-documented ACM mouse models^32–38^. The advantage of this approach, compared with gene supplementation therapies aimed at reintroducing functional gene copies of the defective causal gene, is that it may be applicable to genetic lesions in several different genes.

It now seems essential to consider the broader notion of an adhesome, where lateral membrane signaling participates in the remodeling of the ID. This insight could lead to a better understanding of the mechanisms of ACM and open up new avenues for the potential treatment of this genetic pathology for which no curative treatment currently exists.

## ACKNOWLEDGMENTS

The authors acknowledge the excellent advice and carefull editing of the manuscript from Dr. William T. Pu. They also thank the translational vector core (UMR 1089, Nantes University) for the vectors bioproduction, and Morgane Le Gall (PROTEOM’IC, Institut Cochin, Paris) for IPA analysis, and Dr. Alain Schmitt (Paris Cité University, PIME core facility, Institu Cochin, Paris).

## REFERENCES

1. Balse E, Steele DF, Abriel H, Coulombe A, Fedida D, Hatem SN. Dynamic of ion channel expression at the plasma membrane of cardiomyocytes. Physiol Rev. 2012;92:1317–1358.

2. Austin KM, Trembley MA, Chandler SF, Sanders SP, Saffitz JE, Abrams DJ, Pu WT. Molecular mechanisms of arrhythmogenic cardiomyopathy. Nat Rev Cardiol. 2019;16:519–537.

3. Corrado D, Link MS, Calkins H. Arrhythmogenic Right Ventricular Cardiomyopathy. N Engl J Med. 2017;376:61–72.

4. Towbin JA. Inherited cardiomyopathies. Circ J. 2014;78:2347–2356.

5. Gandjbakhch E, Redheuil A, Pousset F, Charron P, Frank R. Clinical Diagnosis, Imaging, and Genetics of Arrhythmogenic Right Ventricular Cardiomyopathy/Dysplasia: JACC State-of-the-Art Review. J Am Coll Cardiol. 2018;72:784–804.

6. Funke L, Dakoji S, Bredt DS. Membrane-associated guanylate kinases regulate adhesion and plasticity at cell junctions. Annu Rev Biochem. 2005;74:219–245.

7. Montgomery JM, Zamorano PL, Garner CC. MAGUKs in synapse assembly and function: an emerging view. Cell Mol Life Sci. 2004;61:911–929.

8. Hsueh Y-P. The role of the MAGUK protein CASK in neural development and synaptic function. Curr Med Chem. 2006;13:1915–1927.

9. Eichel CA, Beuriot A, Chevalier MYE, Rougier J-S, Louault F, Dilanian G, Amour J, Coulombe A, Abriel H, Hatem SN, Balse E. Lateral Membrane-Specific MAGUK CASK Down-Regulates NaV1.5 Channel in Cardiac Myocytes. Circ Res. 2016;119:544–556.

10. Beuriot A, Eichel CA, Dilanian G, Louault F, Melgari D, Doisne N, Coulombe A, Hatem SN, Balse E. Distinct calcium/calmodulin-dependent serine protein kinase domains control cardiac sodium channel membrane expression and focal adhesion anchoring. Heart Rhythm. 2020;17:786–794.

11. Vite A, Gandjbakhch E, Hery T, Fressart V, Gary F, Simon F, Varnous S, Hidden Lucet F, Charron P, Villard E. Desmoglein-2 mutations in propeptide cleavage-site causes arrhythmogenic right ventricular cardiomyopathy/dysplasia by impairing extracellular 1-dependent desmosomal interactions upon cellular stress. Europace. 2020;22:320–329.

12. Boycott HE, Barbier CSM, Eichel CA, Costa KD, Martins RP, Louault F, Dilanian G, Coulombe A, Hatem SN, Balse E. Shear stress triggers insertion of voltage-gated potassium channels from intracellular compartments in atrial myocytes. Proc Natl Acad Sci U S A. 2013;110:E3955–3964.

13. Pierre B, Laëtitia D-B, Camille B, Claire P, Elise B, Estelle G, Vincent F, Eric V. Generation of CRISPR/Cas9 edited human induced pluripotent stem cell line carrying the heterozygous p.H695VfsX5 frameshift mutation in the exon 10 of the PKP2 gene. Stem Cell Res. 2024;76:103341.

14. Ader F, Duboscq-Bidot L, Marteau S, Hamlin M, Richard P, Fontaine V, Villard E. Generation of CRISPR-Cas9 edited human induced pluripotent stem cell line carrying FLNC exon skipping variant. Stem Cell Res. 2022;58:102616.

15. Novak P, Li C, Shevchuk AI, Stepanyan R, Caldwell M, Hughes S, Smart TG, Gorelik J, Ostanin VP, Lab MJ, Moss GWJ, Frolenkov GI, Klenerman D, Korchev YE. Nanoscale live-cell imaging using hopping probe ion conductance microscopy. Nat Methods. 2009;6:279–281.

16. Gorelkin P, Erofeev A, Kolmogorov V, Efremov Y, Novak P, Shevchuk A, Majouga A, Korchev Y. Scanning Ion Conductance Microscopy (SICM) for Low-stress Directly Examining of Cellular Mechanics. Microscopy and Microanalysis. 2020;26:1968–1970.

17. Clarke RW, Novak P, Zhukov A, Tyler EJ, Cano-Jaimez M, Drews A, Richards O, Volynski K, Bishop C, Klenerman D. Low Stress Ion Conductance Microscopy of Sub-Cellular Stiffness †Electronic supplementary information (ESI) available: Supplementary methods and expanded data presentations of approach curves, stiffness maps, and nanopipet stresses and forces. See DOI: 10.1039/c6sm01106c Click here for additional data file. Click here for additional data file. Soft Matter. 2016;12:7953–7958.

18. Verkerk AO, van Ginneken ACG, van Veen TAB, Tan HL. Effects of heart failure on brain-type Na+ channels in rabbit ventricular myocytes. Europace. 2007;9:571–577.

19. Lin X, Liu N, Lu J, Zhang J, Anumonwo JMB, Isom LL, Fishman GI, Delmar M. Subcellular heterogeneity of sodium current properties in adult cardiac ventricular myocytes. Heart Rhythm. 2011;8:1923–1930.

20. Swiatlowska P, Sanchez-Alonso JL, Wright PT, Novak P, Gorelik J. Microtubules regulate cardiomyocyte transversal Young’s modulus. Proc Natl Acad Sci U S A. 2020;117:2764–2766.

21. Etienne-Manneville S, Hall A. Rho GTPases in cell biology. Nature. 2002;420:629–635.

22. Discher DE, Janmey P, Wang Y-L. Tissue cells feel and respond to the stiffness of their substrate. Science. 2005;310:1139–1143.

23. Kothari P, Johnson C, Sandone C, Iglesias PA, Robinson DN. How the mechanobiome drives cell behavior, viewed through the lens of control theory. J Cell Sci. 2019;132:jcs234476.

24. Kaplan SR, Gard JJ, Protonotarios N, Tsatsopoulou A, Spiliopoulou C, Anastasakis A, Squarcioni CP, McKenna WJ, Thiene G, Basso C, Brousse N, Fontaine G, Saffitz JE. Remodeling of myocyte gap junctions in arrhythmogenic right ventricular cardiomyopathy due to a deletion in plakoglobin (Naxos disease). Heart Rhythm. 2004;1:3–11.

25. Rasmussen TB, Nissen PH, Palmfeldt J, Gehmlich K, Dalager S, Jensen UB, Kim WY, Heickendorff L, Mølgaard H, Jensen HK, Baandrup UT, Bross P, Mogensen J. Truncating plakophilin-2 mutations in arrhythmogenic cardiomyopathy are associated with protein haploinsufficiency in both myocardium and epidermis. Circ Cardiovasc Genet. 2014;7:230–240.

26. Saffitz JE, Asimaki A, Huang H. Arrhythmogenic right ventricular cardiomyopathy: new insights into disease mechanisms and diagnosis. J Investig Med. 2009;57:861–864.

27. Tandri H, Asimaki A, Dalal D, Saffitz JE, Halushka MK, Calkins H. Gap junction remodeling in a case of arrhythmogenic right ventricular dysplasia due to plakophilin-2 mutation. J Cardiovasc Electrophysiol. 2008;19:1212–1214.

28. Vite A, Gandjbakhch E, Prost C, Fressart V, Fouret P, Neyroud N, Gary F, Donal E, Varnous S, Fontaine G, Fornes P, Hidden-Lucet F, Komajda M, Charron P, Villard E. Desmosomal cadherins are decreased in explanted arrhythmogenic right ventricular dysplasia/cardiomyopathy patient hearts. PLoS One. 2013;8:e75082.

29. Noorman M, Hakim S, Kessler E, Groeneweg JA, Cox MGPJ, Asimaki A, van Rijen HVM, van Stuijvenberg L, Chkourko H, van der Heyden MAG, Vos MA, de Jonge N, van der Smagt JJ, Dooijes D, Vink A, de Weger RA, Varro A, de Bakker JMT, Saffitz JE, Hund TJ, Mohler PJ, Delmar M, Hauer RNW, van Veen TAB. Remodeling of the cardiac sodium channel, connexin43, and plakoglobin at the intercalated disk in patients with arrhythmogenic cardiomyopathy. Heart Rhythm. 2013;10:412–419.

30. Tsui H, van Kampen SJ, Han SJ, Meraviglia V, van Ham WB, Casini S, van der Kraak P, Vink A, Yin X, Mayr M, Bossu A, Marchal GA, Monshouwer-Kloots J, Eding J, Versteeg D, de Ruiter H, Bezstarosti K, Groeneweg J, Klaasen SJ, van Laake LW, Demmers JAA, Kops GJPL, Mummery CL, van Veen TAB, Remme CA, Bellin M, van Rooij E. Desmosomal protein degradation as an underlying cause of arrhythmogenic cardiomyopathy. Sci Transl Med. 2023;15:eadd4248.

31. Asimaki A, Kapoor S, Plovie E, Karin Arndt A, Adams E, Liu Z, James CA, Judge DP, Calkins H, Churko J, Wu JC, MacRae CA, Kléber AG, Saffitz JE. Identification of a new modulator of the intercalated disc in a zebrafish model of arrhythmogenic cardiomyopathy. Sci Transl Med. 2014;6:240ra74.

32. Cerrone M, Lin X, Zhang M, Agullo-Pascual E, Pfenniger A, Chkourko Gusky H, Novelli V, Kim C, Tirasawadichai T, Judge DP, Rothenberg E, Chen H-SV, Napolitano C, Priori SG, Delmar M. Missense mutations in plakophilin-2 cause sodium current deficit and associate with a Brugada syndrome phenotype. Circulation. 2014;129:1092–1103.

33. Moncayo-Arlandi J, Guasch E, Sanz-de la Garza M, Casado M, Garcia NA, Mont L, Sitges M, Knöll R, Buyandelger B, Campuzano O, Diez-Juan A, Brugada R. Molecular disturbance underlies to arrhythmogenic cardiomyopathy induced by transgene content, age and exercise in a truncated PKP2 mouse model. Hum Mol Genet. 2016;25:3676–3688.

34. van Opbergen CJM, Noorman M, Pfenniger A, Copier JS, Vermij SH, Li Z, van der Nagel R, Zhang M, de Bakker JMT, Glass AM, Mohler PJ, Taffet SM, Vos MA, van Rijen HVM, Delmar M, van Veen TAB. Plakophilin-2 Haploinsufficiency Causes Calcium Handling Deficits and Modulates the Cardiac Response Towards Stress. Int J Mol Sci. 2019;20:4076.

35. Camors EM, Roth AH, Alef JR, Sullivan RD, Johnson JN, Purevjav E, Towbin JA. Progressive Reduction in Right Ventricular Contractile Function Attributable to Altered Actin Expression in an Aging Mouse Model of Arrhythmogenic Cardiomyopathy. Circulation. 2022;145:1609–1624.

36. Bradford WH, Zhang J, Gutierrez-Lara EJ, Liang Y, Do A, Wang T-M, Nguyen L, Mataraarachchi N, Wang J, Gu Y, McCulloch A, Peterson KL, Sheikh F. Plakophilin 2 gene therapy prevents and rescues arrhythmogenic right ventricular cardiomyopathy in a mouse model harboring patient genetics. Nat Cardiovasc Res. 2023;2:1246–1261.

37. Fabritz L, Fortmueller L, Gehmlich K, Kant S, Kemper M, Kucerova D, Syeda F, Faber C, Leube RE, Kirchhof P, Krusche CA. Endurance Training Provokes Arrhythmogenic Right Ventricular Cardiomyopathy Phenotype in Heterozygous Desmoglein-2 Mutants: Alleviation by Preload Reduction. Biomedicines. 2024;12:985.

38. Rizzo S, Lodder EM, Verkerk AO, Wolswinkel R, Beekman L, Pilichou K, Basso C, Remme CA, Thiene G, Bezzina CR. Intercalated disc abnormalities, reduced Na(+) current density, and conduction slowing in desmoglein-2 mutant mice prior to cardiomyopathic changes. Cardiovasc Res. 2012;95:409–418.

