## Supplemental Figures and Tables for "Depleting trafficking regulator CASK promotes intercalated disc organization and ventricular function"

### SUPPLEMENTAL FIGURE 1

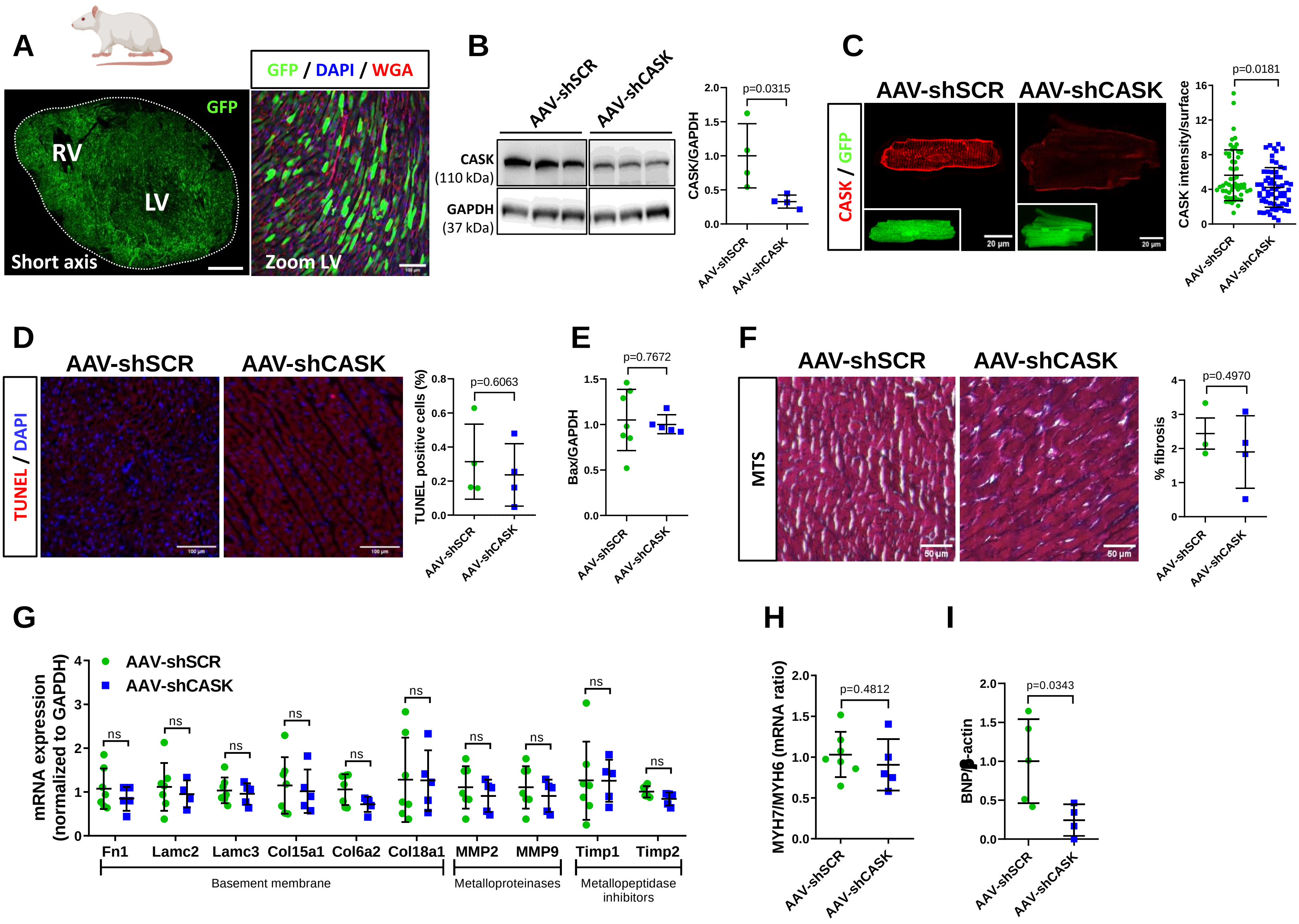

### SUPPLEMENTAL FIGURE 2

**A**

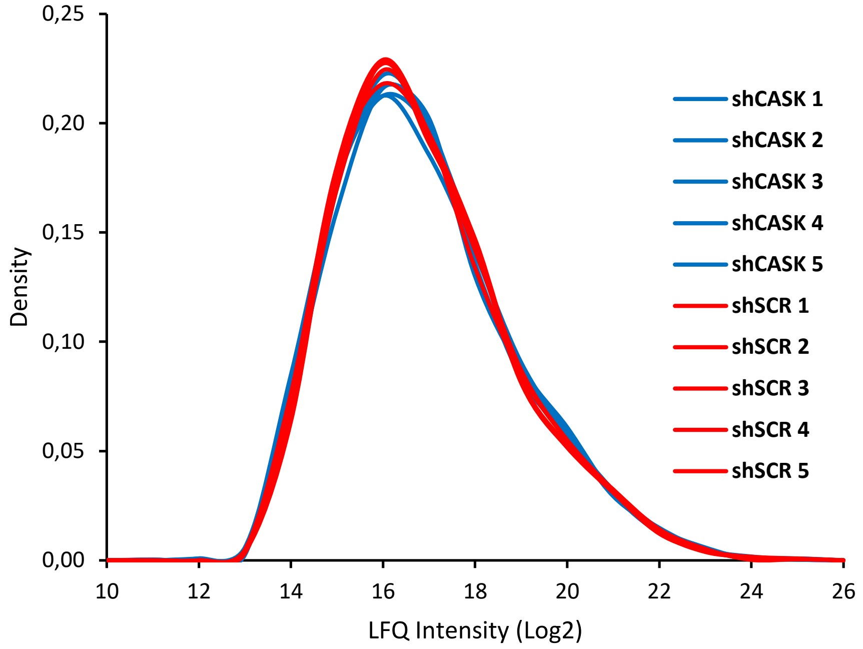


**B**

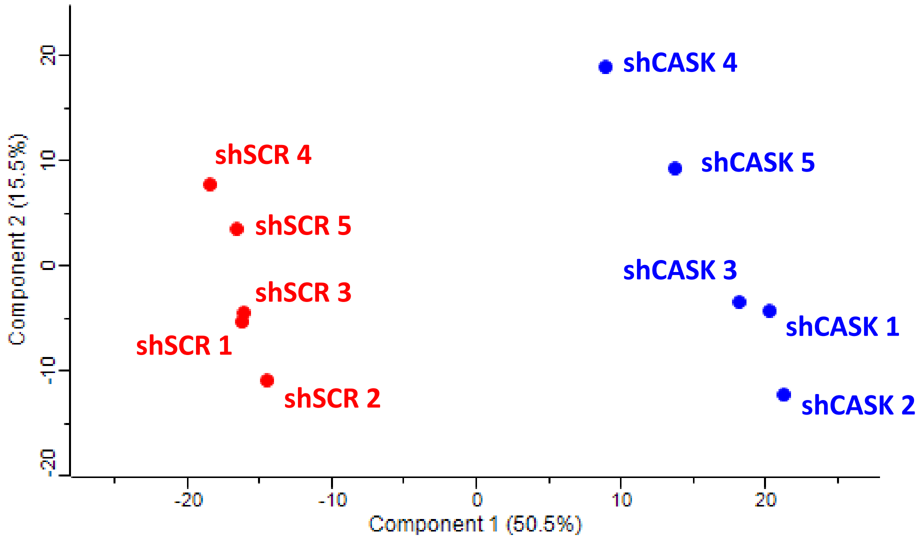

### SUPPLEMENTAL FIGURE 3

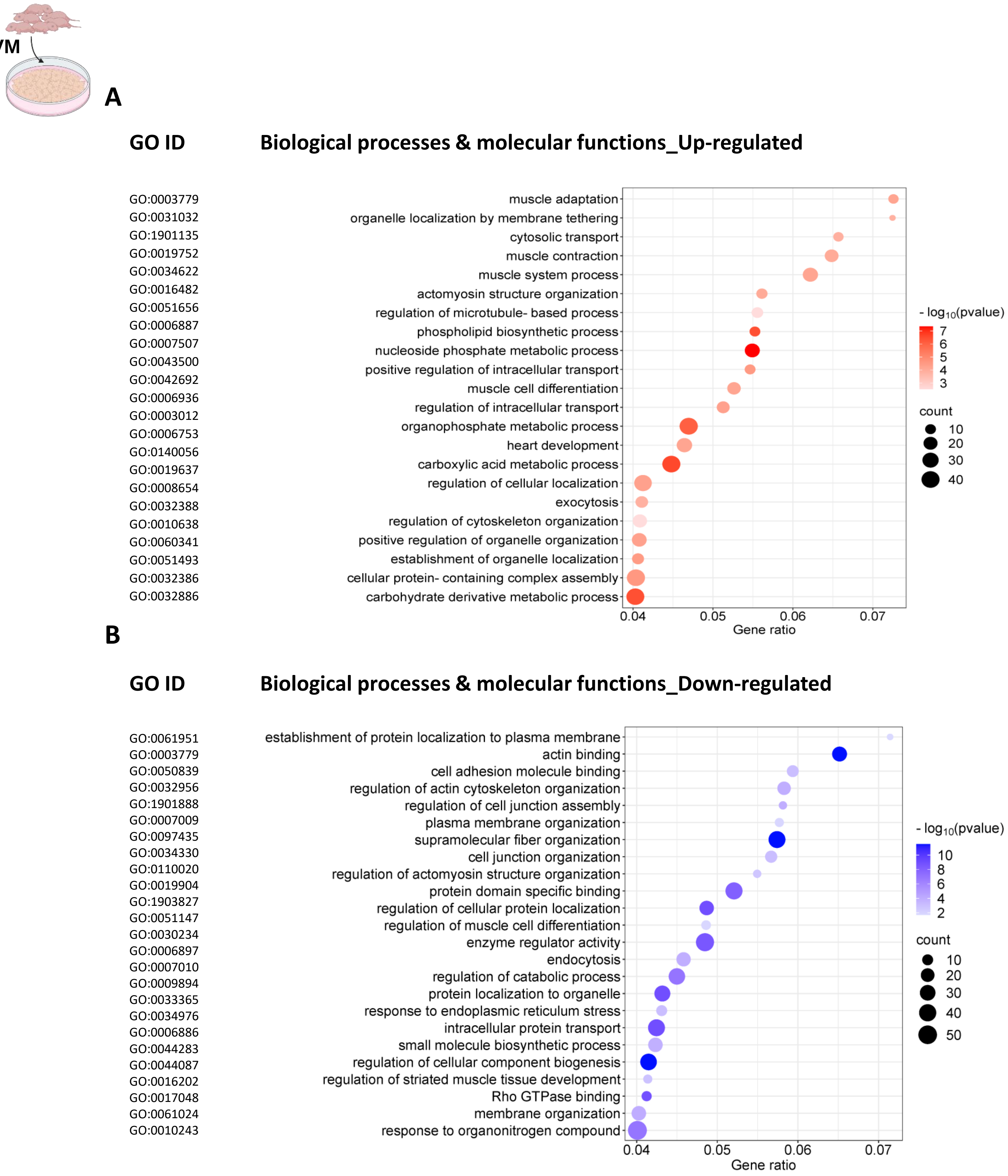

### SUPPLEMENTAL FIGURE 4

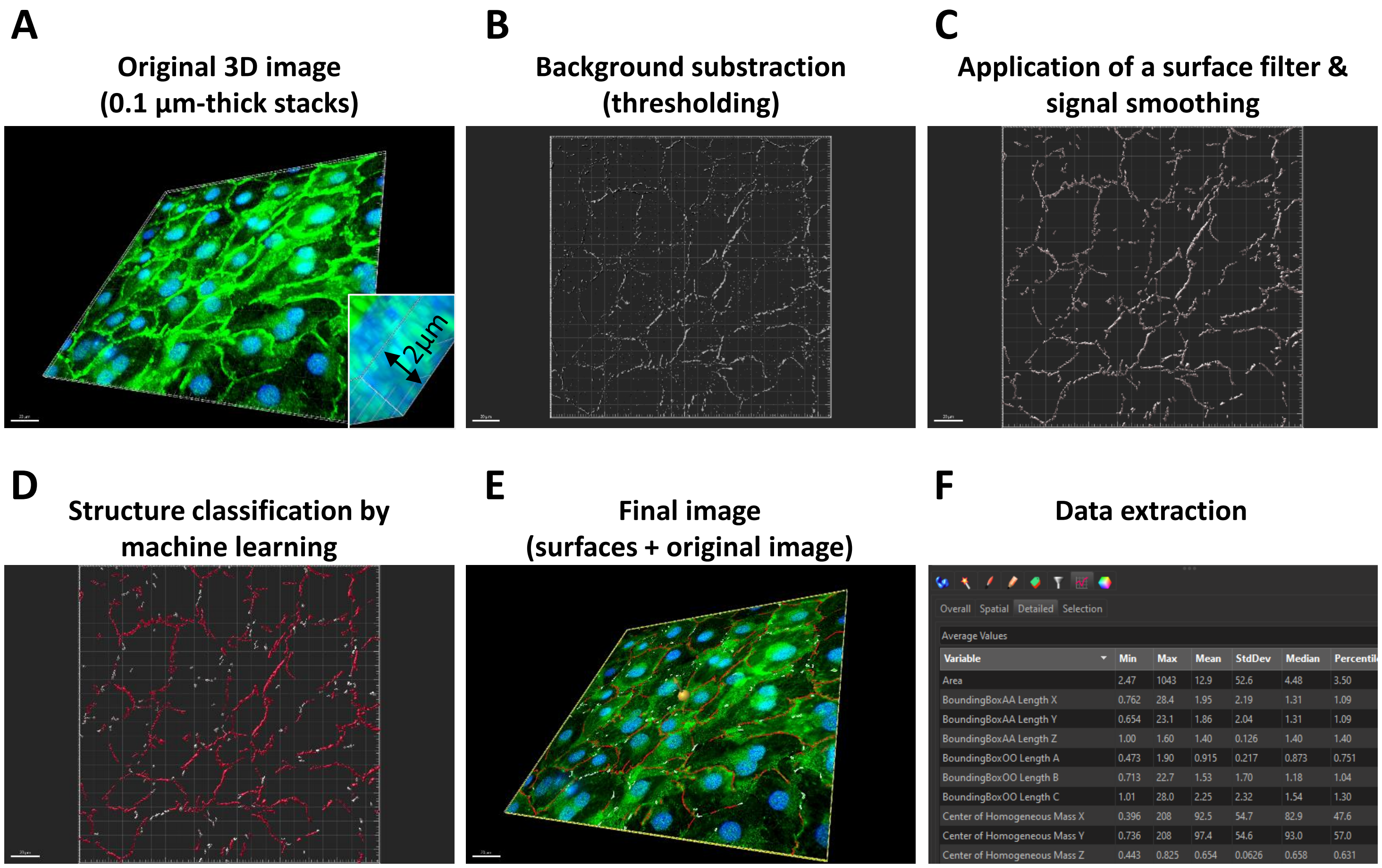

### SUPPLEMENTAL FIGURE 5

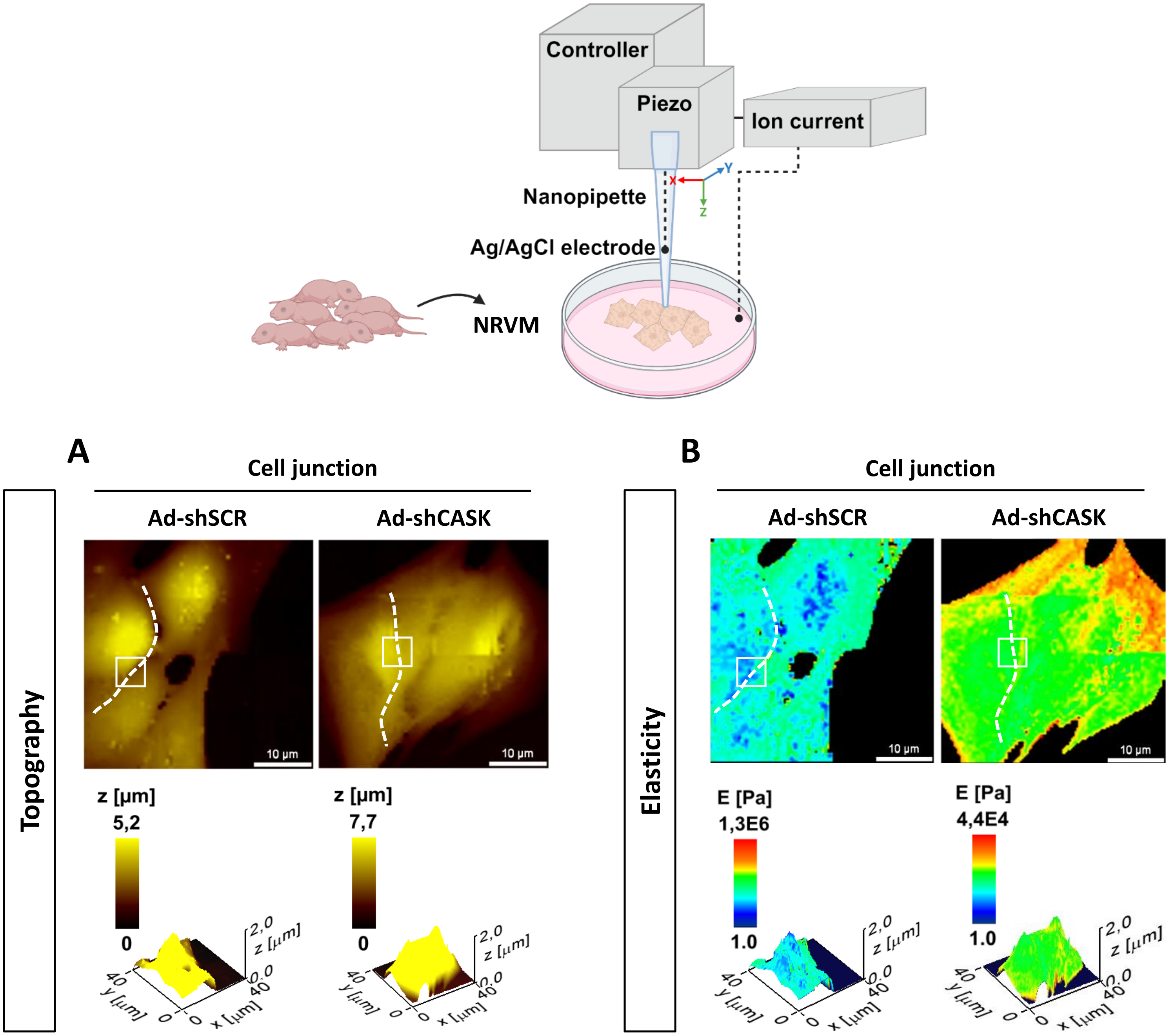

### SUPPLEMENTAL FIGURE 6

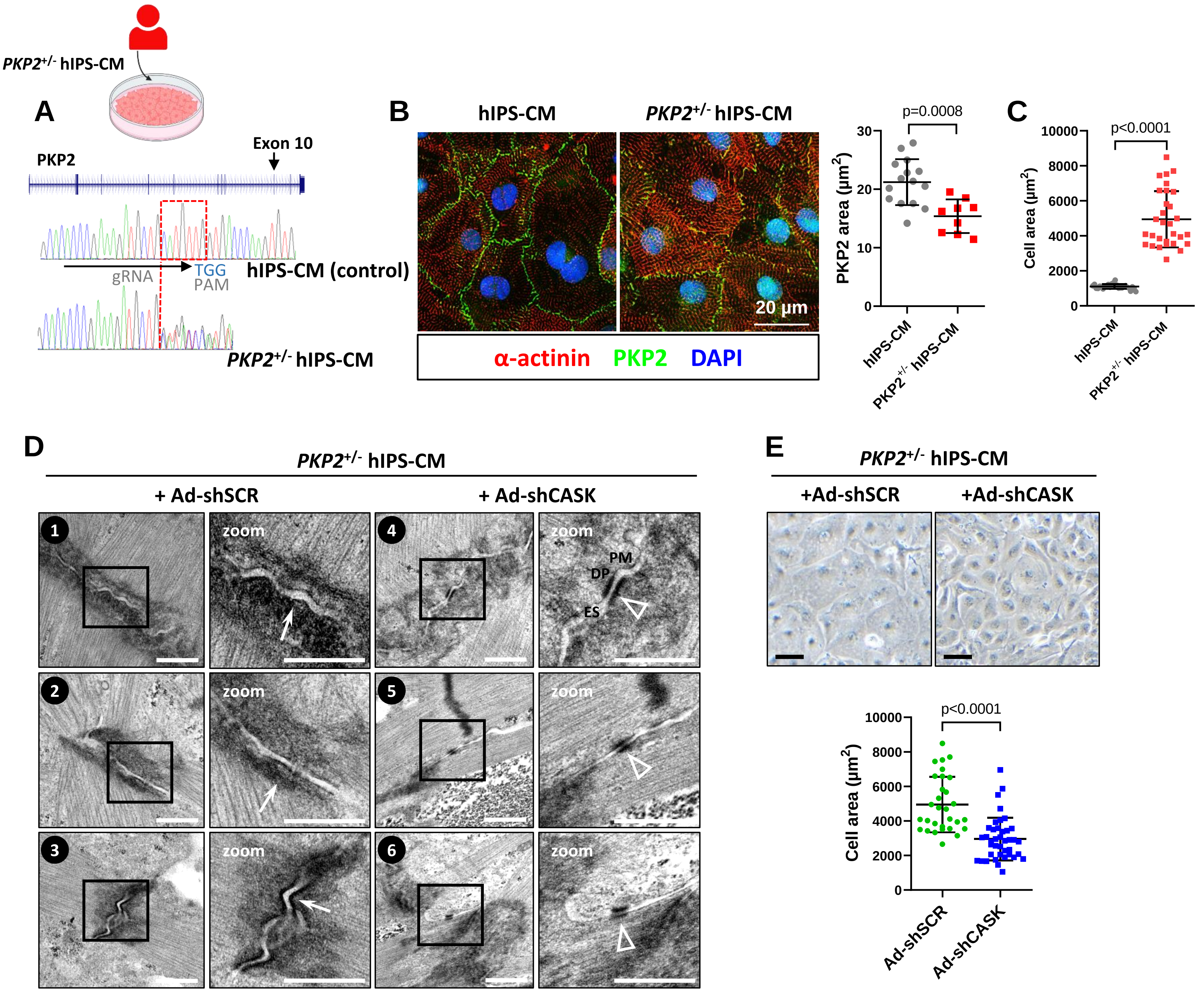

### SUPPLEMENTAL TABLE 1

## A

| | Echograhy 60 days post-AAV | |
|---|---|---|
| | **AAV-shSCR** | **AAV-shCASK** |
| ***N*** | *23* | *21* |
| ***End-diastolic parameters (mm)*** | | |
| IVS | 1.33 ± 0.04 | 1.2 ± 0.03* |
| LVPW | 1.53 ± 0.04 | 1.26 ± 0.03**** |
| LVED | 6.43 ± 0.11 | 7.59 ± 0.14**** |
| ***End-systolic parameters (mm)*** | | |
| IVS | 2.34 ± 0.05 | 1.98 ± 0.04**** |
| LVPW | 2.19 ± 0.05 | 1.98 ± 0.04* |
| LVED | 3.29 ± 0.09 | 4.68 ± 0.12**** |
| ***Remodeling index*** | | |
| Relative wall thickness | 0.45 ± 0.04 | 0.32 ± 0.04**** |
| ***Left ventricular function*** | | |
| LVEF (%) | 85.06 ± 0.56 | 74.29 ± 0.89**** |
| LVFS (%) | 49.03 ± 0.72 | 38.63 ± 0.76**** |
| Stroke volume (mL) | 0.53 ± 0.12 | 0.73 ± 0.14**** |
| Heart rate (BPM) | 387.70 ± 50.38 | 354.00 ± 35.94* |
| Cardiac output (mL/min) | 202.60 ± 41.95 | 256.60 ± 51.54*** |

## B

| | ECG 60 days post-AAV | |
|---|---|---|
| | **AAV-shSCR** | **AAV-shCASK** |
| ***N*** | *18* | *20* |
| **RR (ms)** | 154.4±17.5 | 138.1±16.6 |
| **HR (BPM)** | 418.9±51.1 | 441.5±53.2 |
| **PR (ms)** | 38.6±4.5 | 38.0±3.3 |
| **Pdur (ms)** | 21.0±4.5 | 22.05±3.4 |
| **QRS (ms)** | 37.4±6.6 | 39.7±7.2 |
| **QT (ms)** | 72.0±5.8 | 75.1±8.9 |
| **QTcB (ms)** | 189.7±14.6 | 202.6±20.9 |

## C

| | Flecainide challenge P60 days post-AAV injection | | | |
|---|---|---|---|---|
| | **AAV-shSCR** | **AAV-shSCR** | **AAV-shCASK** | **AAV-shCASK** |
| | **- Flecainide** | **+ Flecainide** | **- Flecainide** | **+ Flecainide** |
| ***N*** | *7* | *7* | *8* | *8* |
| **RR (ms)** | 140.7±19.3 | 149.6±24.9 | 132.1±16.8 | 143.9±14.8 |
| **HR (BPM)** | 434.4±58.9 | 411.1±67.7 | 461.6±56.4 | 420.9±40.9 |
| **PR (ms)** | 38.35±5.1 | 40.6±5.7 | 38.8±2.7 | 42.3±6.5 |
| **Pdur (ms)** | 20.1±1.8 | 20.3±1.4 | 20.1±1.9 | 19.8±1.3 |
| **QRS (ms)** | 37.1±7.8 | 35.9±9.4 | 38.1±8.2 | 39.0±9.9 |
| **QT (ms)** | 72.8±7.1 | 75.4±13.5 | 73.8±11.9 | 77.8±10.8 |
| **QTcB (ms)** | 194.7±11.6 | 195.9±29.8 | 202.8±25.4 | 206.2±32.3 |

### SUPPLEMENTAL TABLE 2

| ARVC | Desmosomal mutation | Sex | Age at diagnosis | Age at HtTx | Arrhythmias | Morphology | Histology | Use |
|---|---|---|---|---|---|---|---|---|
| 1 | DGS2 (c.690+1 G>A) | M | 50 | 73 | Multifocal VT from RV and LV | Major RV dilation and hypokinesia | Extensive fibro-fatty replacement | WB |
| 2 | DSG2 (p.Arg49His) | M | 13 | 14 | Multifocal VT from RV | Major RV dilation and hypokinesia | Extensive sub-epicardial fibro-fatty replacement | IHC |
| 4 | None | F | 21 | 30 | Multifocal VT from RV | Major RV dilation and hypokinesia | Extensive fibro-fatty replacement | WB |
| 3 | None | M | 46 | 53 | Multifocal VT from RV, FV | Major RV dilation, anterior akinesia, severe hypokinesia | Extensive fibro-fatty replacement | WB |
| 6 | DSG2 (p.Arg46Trp) | M | 39 | 48 | Multifocal VT from RV | Major RV dilation and hypokinesia | Extensive fibro-fatty replacement | WB, IHC |
| 8 | PKP2 | | | | | | | WB |
